# Enhancing Lipid Production in *Yarrowia lipolytica* through Continuous Fermentation and Adaptive Laboratory Evolution in Bioreactors

**DOI:** 10.1101/2025.08.20.671336

**Authors:** Elif Kurt, Lucas Sa, Evelyn Anjo, Dongming Xie

**Author notes:** Corresponding Author: Dr. Dongming Xie.

## Abstract

*Yarrowia lipolytica* excels in microbial lipid production, thriving across diverse conditions. Batch or fed-batch fermentation is the common practice to achieve higher lipid titer and yield, but it is also subject to lower lipid productivity. Single-stage continuous fermentation (CF) provides a great potential for significantly higher productivity, but genetic instability is often seen and challenges strain performance over the long-period CF. This study harnesses single-stage CF to not only improve lipid productivity, but also evolve high-lipid mutants from a previously engineered *Y. lipolytica* strain E26 via adaptive laboratory evolution (ALE) in a continuous bioreactor, guided by a predictive kinetic model. The single-stage CF was run for 1128 hours (47 days) with key process parameter adjusted in a 1-L bioreactor to produce over 150 g/L yeast biomass, exceeding the targeted 113 g/L that is predicted by the model. Compared with the fed-batch fermentation process, the single-stage CF successfully improved lipid productivity from 0.3∼0.5 g/L/h to about 1 g/L/h while maintained the lipid yield at around 0.1 g/g. The CF sample at 1008 h was used to isolate mutants with higher lipid production after ALE in the continuous bioreactor. A mutant E26E03 was identified which demonstrated improvements in biomass, lipid content, and lipid yield by 43%, 30%, and 51%, respectively, over the original strain E26 in fed-batch fermentation. Our study indicated that using model-guided CF with ALE in a continuous bioreactor provides a great potential for significantly higher product titer, rate, and yield (TRY) in biomanufacturing.

## 1. Introduction

*Yarrowia lipolytica* is one of the most promising microbes for sustainable lipid production, offering renewable alternative to plant- and animal-derived oils. This yeast excels in accumulating lipids up to 80% of its dry cell weight (DCW) under nitrogen-limited conditions and can utilize diverse carbon sources^1^. Recent advances have further expanded its potential, with engineered strains achieving tailored lipid profiles such as omega-3 fatty acids^2^. Despite these strengths, scaling *Y. lipolytica* production is hampered by limitations in traditional fermentation systems. Batch fermentation systems are constrained by substrate inhibition, toxic byproduct accumulation, and low productivity, leading to suboptimal titer, rate, and yield (TRY) metrics^3,4^. Fed-batch setup alleviates some issues but require complex feeding strategies to maintain optimal carbon-to-nitrogen (C/N) ratios, often resulting in inconsistent yields and prolonged operational downtime^5^.

Continuous fermentation (CF) offers a promising alternative by enabling steady-state operation, higher biomass densities, and reduced process complexity, as demonstrated in recent studies achieving stable lipid production over extended runs^6^. However, prolonged CF runs risk genetic instability, leading to reduced lipid accumulation and strain performance over time^7^. Adaptive laboratory evolution (ALE) provides a powerful approach to address genetic instability by selecting mutants with enhanced lipid production, stress tolerance, and process robustness under continuous culture conditions^8,9^. Recent ALE studies with *Y. lipolytica* have identified mutants with improved lipid/DCW ratios and tolerance to low pH or high glucose^10,11^, critical for industrial scalability^12^. Kinetic modeling further enhances CF by predicting optimal parameters, such as dilution rates and dissolved oxygen (DO) setpoints^13,14^. These advancements underscore the potential of integrating CF with ALE and modeling to overcome bioprocess limitations.

This study aims to develop a model-guided CF platform coupled with ALE to evolve high-lipid-producing mutants of *Y. lipolytica* E26^15^, a strain engineered for lipid overproduction. By optimizing key parameters and reinforcing long-term cultivation to select robust mutants, this work aims to enhance TRY of lipid production and isolate an enhanced lipid producer by a new ALE platform in continuous bioreactor. The kinetic model will also be employed and tailored to guide the design of continuous feed and dilution rate and used as a performance checker throughout the long-term CF.

## 2. Materials and methods

### 2.1. Strains and media

*Yarrowia lipolytica* strain used in all experiments and subjected to further adaptive laboratory evolution, E26, was obtained from Dr. Alper’s group, which was generated via overexpression of the type I diacylglycerol acyltransferase and disruption of β-oxidation in *Y. lipolytica* Po1f (PO1f *Δpex10Δmfe* leucine+ uracil+ DGA1)^15^.

The seed culture medium was composed of 5 g/L yeast extract (VWR, USA), 5 g/L ammonium sulfate (Fisher Scientific, USA), 6 g/L potassium phosphate monobasic (Fisher Scientific, USA), 2 g/L sodium phosphate dibasic (Thermo Fisher, USA), and 40 g/L D-glucose (VWR, USA).

The initial fermentation media was the same for both fed-batch and CF and contained 12 g/L yeast extract, 9 g/L ammonium sulfate, 6 g/L potassium phosphate monobasic, 2 g/L sodium phosphate dibasic, 50 g/L D-glucose, 1.5 mg/L Thiamin·HCl (Thermo Fisher, USA), 2 ml/L trace metals (100X), 0.5 g/L MgSO4, and 1 mL/L Antifoam 204 (Sigma Aldrich, USA). The trace metal (100X) stock solution included 15 g/L citric acid (Fisher Scientific, USA), 1.5 g/L calcium chloride dihydrate (Fisher Scientific, USA), 10 g/L iron sulfate heptahydrate (RPI, USA), 0.39 g/L zinc sulfate heptahydrate (VWR, USA), 0.38 g/L copper sulfate pentahydrate (VWR, USA), and 0.3 g/L manganese sulfate tetrahydrate (VWR, USA). It was filter-sterilized through 0.22mm sterile membrane and stored at -20^°^C.

The feed solution of the CF was reprepared on demand and contained 16 g/L yeast extract 12 g/L ammonium sulfate, 4 g/L potassium phosphate monobasic, 2.5 g/L sodium phosphate dibasic, 0.75 mg/L Thiamin·HCl (0.5 mL/L), 1 mL/L trace metals (100X), 0.25 g/L MgSO4, 100 ug/mL ampicillin and 360∼460 g/L D-glucose (determined by model simulation and further adjustments during the fermentation).

The flask culture medium for colony screening of ALE from continuous bioreactor contained 1.25 g/L yeast extract, 0.6 g/L ammonium sulfate, 4 g/L potassium phosphate monobasic, 13 g/L potassium phosphate dibasic (Fisher Scientific, USA), 0.3 g/L Thiamin·HCl, 0.5 mL/L trace metals (100X), 0.5 g/L MgSO4, and 20 g/L D-glucose.

### 2.2. Fermentation protocols

An overview of the major experimental procedures and protocols for this study is shown in **Figure 1**, which includes fed-batch fermentation, computer model-guided continuous fermentation, off-line sample analysis, mutant screening for high-lipid producers generated through ALE in continuous bioreactor, and further confirmation of top mutant in fed-batch fermentation.

**Figure 1.**
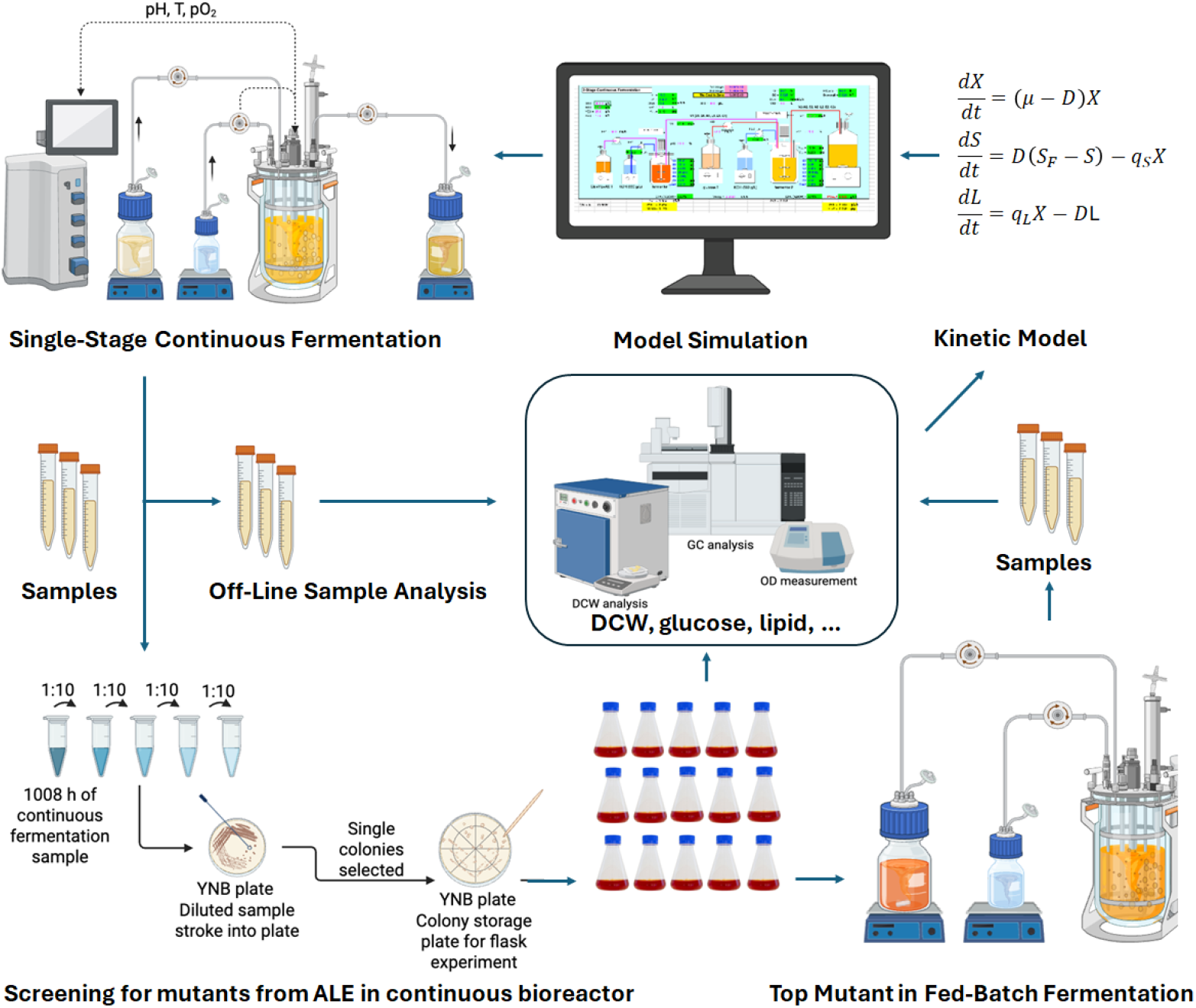
An overview of the major experimental procedures and protocols that were used in this study.

#### 2.2.1. Fed-batch fermentation

Seed vials of *Y. lipolytica* E26 and mutants, which were stored at -80^°^C, were thawed at ambient temperature. To prepare the inoculant, each vial (0.5 mL) was transferred to a 250 mL shake flask containing 50 mL of the seed culture medium described above. Subsequently, these seed cells underwent a growth phase in shake flasks within an Innova-4000 incubator shaker, which was operated at 30^°^C with a shaking speed of 250 rpm for 12∼16 h until the OD600 attained 2∼5.

The shake flask seed culture (50 mL, OD600 = 2∼5) was transferred to a 1-L fermenter (Biostat B-DCU, Sartorius, Germany) to initiate fermentation (t = 0 h). The initial working volume was 0.7 L. Glucose (50 g/L) was used as the initial substrate. The pO_2_ level of the fermentation experiments was set at 20% air saturation by cascade controls of agitation speed between 500 and 1200 rpm, unless otherwise specified, and pure oxygen enrichment if needed. The aeration rate was fixed at 0.5 L/min or otherwise specified. The temperature was maintained at 30 ^°^C throughout the experiments. The pH value was maintained at 3.5∼7.0 based on each specific experiment design by feeding KOH (10 M). Glucose feeding (600 g/L) commenced when the initial glucose concentration dropped below 15 g/L and it was maintained at approximately 10∼20 g/L by adjusting the glucose feed rate based on offline glucose measurements. Samples (∼10 mL) were taken every 6∼12 h throughout the run for off-line analysis of cell density (OD600, DCW), residual glucose, and lipid content in yeast biomass.

#### 2.2.2. Single-stage continuous fermentation

The CF of *Y. lipolytica* E26 was conducted in a 1-L bioreactor containing 0.7-L working volume. The initial CF medium and temperature were the same as described in the fed-batch fermentation protocols. The initial pH was set at 6.0; the dissolved oxygen (DO, or represented by pO_2_) was set at 20% of air saturation; and the targeted residual glucose concentration was maintained at 10-15 g/L. The initial feed for CF of *Y. lipolytica* E26 was predetermined by model simulation, as described in more detail in Kinetic Modeling and Supplementary Information, which included 16 g/L yeast extract, 12 g/L (NH_4_)_2_SO_4_, and 360 g/L glucose. The feed rate was preset at ∼17 mL/L to result in a dilution rate of ∼0.025 h^-1^ and a residence time of ∼40 h. The feeding started when the initial glucose in the medium dropped below 15 g/L and it was slightly adjusted to maintain glucose within 10∼20 g/L during the CF process. At the same time, a steel tube for sampling, connected with an external fast-running pump, was installed in the bioreactor with its lower end touching only the liquid culture surface to allow for continuous broth bleeding and maintain a constant working volume of 0.7 L.

During the CF experiment, several process parameters, including the pH value, DO, and glucose concentration in feed, were changed to test their effects on cell growth and lipid production for further optimization. Typically a new change of the process parameter(s) applied only ∼120 h after the previous change to allow the CF reach a relatively steady state, which requires a period of 3-fold residence time or longer. The changes included (1) increasing the pH value from 6.0 to 7.0 at 216 h, (2) lowering the PH value from 7.0 to 5.0 at 360 h, (3) increasing the glucose concentration in feed from 360 g/L to 460 g/L at 456 h and adjusting the feeding rate accordingly to maintain the glucose concentration within 10∼20 g/L in fermentation medium, (4) decreasing DO from 20% to 10% air saturation at 744 h, (5) adjusting all parameters back to pH 5.0, DO 20%, and 360 g/L glucose in feed at 960 h, (6) lowering the pH value from 5.0 to 3.5 at 1080 h and maintaining all other parameters till the end of the CF experiment. Samples (∼10 mL) were taken every 12∼24 h throughout the run for off-line analysis of cell density (OD600, DCW), residual glucose, lipid content in yeast biomass, and colony screening for potential mutants with enhanced lipid production capability.

### 2.3. Colony screening for mutants from ALE in continuous bioreactor

The fermentation sample from the timepoints at 1008 h was used for colony screening for mutants from ALE in the continuous bioreactor. About 0.1 mL of the sample was diluted by 10^7^ times and stroked onto a YNB (6.7 g/L yeast nitrogen base + 20 g/L glucose) plate, which was incubated at 30^°^C for 2∼3 days until significant growth of single colonies (with ∼1 mm size) was observed. A total of 15 single colonies were selected from the YNB plate and re-stroke onto a new YNB plate for storage. Then, each single colonies was inoculated into 25 mL media in a 125-mL baffled flask, which was cultivated in an Innova-4000 incubator shaker at 30^°^C with a shaking speed of 250 rpm for up to 2 days until the OD_600nm_ reached about 2. About 0.5 mL cell culture sample was taken from each flask culture and mixed with 0.5 mL sterilized glycerol stock solution (50%, w/w) to prepare glycerol stock seed vials, which was stored in a -80^°^C freezer for further investigation.

To prepare the flask cultures for mutant screening, 0.5 mL seed vial solution was used to inoculate 25 mL fed-batch flask culture medium. The flask cultures were conducted in an Innova-4000 incubator shaker at 30^°^C with a shaking speed of 250 rpm for up to 6 days. Two doses of glucose feed (containing 300 g/L glucose) was added into the flask culture at 48 h and 96 h, respectively, to control the residual glucose concentrations within 10∼50 g/L based on the off-line glucose measurements. About 10 mL culture sample from the end of each flask experiment was taken for off-line analysis of cell density (OD600, DCW), residual glucose, and lipid content in yeast biomass. The colonies led to the highest lipid production in the flask cultures were selected for further investigation in fed-batch fermentation to verify the selected mutants’ performance.

### 2.4. Analytical methods

#### 2.4.1. Cell density, glucose analysis, and citric acid formation

Cell density was determined by measuring the optical density at 600 nm (OD600) using a Genesys 10 Vis spectrophotometer (Thermo Fisher Scientific, USA). To measure the dry cell weight (DCW), cells from 2∼5 mL of culture were collected into a 15 mL tube, centrifuged at 3000 rpm for 10 min and washed twice with 5 mL distilled water. Harvested cells were resuspended and dried in a pre-weighed aluminum tray for ∼48 h at 75 ^°^C incubator until a constant weight was reached and then weighed to calculate the DCW. To measure the glucose concentrations in fermentation samples, cell broth in 1.5-mL microtubes was centrifuged for 5 min, then 0.1 mL supernatant was diluted by 10∼100 fold to control the diluted sample’s glucose concentration within 0.1∼2.5 g/L. Glucose levels were measured using a YSI 2950 biochemical analyzer (YSI Inc., USA). The concentration of citric acid was measured with the EnzyChrom Citrate Assay Kit (VWR) as described previously^16^. For fermentation with pH values controlled within 5∼7, the produced citric acid after 24 h was also estimated by the total base KOH (8.5 M) used for pH adjustment.

#### 2.4.2. Lipid analysis

##### 2.4.2.1. TAG and FFA analysis

Lipids synthesized by *Y. lipolytica* were quantified using gas chromatography coupled to a flame ionization detector (GC-FID). Prior to GC analysis, intracellular and extracellular triacylglycerides (TAGs) and free fatty acids (FFAs) were converted into their fatty acid methyl ester (FAME) forms. For disrupting the cell membrane and taking intracellular lipids out of the cell to obtain total lipids (TAG or FFA), ∼0.1 mL (or ∼0.3 g) of 0.5-mm glass beads (Scientific Industries, Inc), 0.3 mL toluene, 0.3 mL methanol, 0.27 mL NaCl (1 M), and 0.075 mL of broth were placed in a 2 mL screw cap tube. The tubes were placed on the vortex at maximum speed for 15-20 minutes. After the centrifugation, 0.1 mL of the toluene layer was collected. To quantify intracellular lipids, 0.025 mL of broth was mixed with 1 mL of DI water and centrifuged for 10 min at 10000 rpm. The pellets were collected as waterless as possible by careful pipetting. Cells prepared for GC analysis were further treated using two methods: base-catalyzed esterification for TAG quantification, and acid-catalyzed esterification for TAG + FFA quantification^17,18^. For the base-catalyzed method, 20 μL of 5 g/L triheptadecanoin or tripentadecanoin (Nu-Check Prep, Inc. USA) dissolved in toluene was used as the internal standard. The reaction was conducted by adding 0.5 mL of 2% sodium methoxide (NaOMe) to methanol and incubating at room temperature for 60 min at 250 rpm. After the reaction, 0.2 mL of 1 M NaCl and 0.88 mL of heptane were added. All samples were vortexed thoroughly and centrifuged to separate the top heptane layer containing FAMEs. For the acid-catalyzed method, 20 μL of 5 g/L heptadeconic acid (Nu-Check Prep, Inc. USA) dissolved in toluene was used as the internal standard. The reaction was conducted by adding 2 mL of 9.4% HCl in methanol and incubating in a water bath shaker at 50 ^°^ C for 10 min at 250 rpm. After the reaction, 0.2 mL of 1 M NaCl and 0.88 mL of heptane were added. All samples were vortexed thoroughly and centrifuged to separate the top heptane layer containing FAMEs, which will be used for further analysis with gas chromatographay.

##### 2.4.2.2. Gas chromatography for lipid analysis

Gas chromatography (GC) analysis of the FAMEs was performed using an Agilent 8860 GC equipped with a flame ionization detector (FID). The samples were injected into the GC/FID system equipped with an Agilent DB-FastFAME (20 m) column. The GC was programmed with the following inlet operating parameters: high-purity helium carrier gas set at a constant flow pressure of 35 psi, inlet temperature of 250 ^°^C, and split injection mode with a split ratio of 100. The detector temperature was set at 290 ^°^C, with an airflow rate of 400 mL/min, hydrogen gas flow rate of 40 mL/min, and makeup gas flow rate of 25 mL/min. The GC oven was programmed with the following temperature regime: start at 80 ^°^C, hold for 1 min, ramp to 194 ^°^C at 35 ^°^C/min, hold for 1 min, and ramp to 240 ^°^C at 5 ^°^C/min. Quantification of fatty acids was achieved using standard curves measuring commercial FAME standards purchased from Nu-Check Prep, Inc. (MN, USA). The identification of FAME was achieved by analyzing mixes of each standard in the GC and creating a calibration curve.

### 2.5. Kinetic modeling

To guide the CF experiment, model simulations was used to design the initial feed and dilution rate. A kinetic model tailored to the *Yarrowia lipolytica* strain was derived from the previous fermentation study and modified for prediction of a strain that is capable of producing about 80% lipid/DCW^4^. This model was originally developed to explore the dynamics of the both fed-batch and continuous fermentation for production of general lipid and other functional fatty acids such as omega-3 EPA^4^. The model equations were modified for lipid production via the continuous *Y. lipolytica* fermentation, which are described in more detail in the **Supplementary Information**. The fermentation model includes the kinetic equations for cell growth, glucose consumption, lipid production, citric acid formation, oxygen consumption rate, nitrogen consumption, and fermentation volume change caused by glucose feeding, base feeding, and evaporation. The model was used to simulate a series of CF conditions (e.g. initial medium, DO, oxygen uptake rate (OUR), the feeding rate of glucose, and the concentrations of glucose, yeast extract, ammonium sulfate in the feed) so that an ideal process can be selected for CF experiments to achieve yeast biomass > 100 g DCW/L and lipid content > 60% (g lipid/g DCW) within a maximum OUR of 100 mmol/L/h in the 1-L bioreactor. The selected CF conditions from the simulated scenarios were used to design the initial CF experiment, as described previously.

## 3. Results and discussion

### 3.1. Fed-batch fermentation of *Y. lipolytica* E26

Fed-batch fermentation experiments were first conducted to understand the *Y. lipolytica* strain E26 and the lipid production process. In general, the fermentation starts with a cell growth phase under a nitrogen-rich condition by using yeast extract and/or (NH_4_)_2_SO_4_ as the nitrogen source, followed by a lipid production phase triggered by a nitrogen-limiting or -starving condition when the initial nitrogen source is consumed^4,16,19^. While a significant amount of research has been conducted to optimize the medium and DO level to improve *Y. lipolytica* cell growth and lipid production, very few studies have investigated the effect of the pH value. Therefore, four different pH values (3.5, 5.0, 6.0, 7.0) were tested in this study to examine the effect of pH value on lipid production by the engineered *Y. lipolytica* E26.

As seen in **Figure 2A**, the yeast biomass production was relatively consistent across all tested pH levels. The lipid content in the yeast biomass (Lipid/DCW) peaked at pH 5.0, with the highest value being recorded at 72 h. The pH 3.5 and 6.0 supported moderate lipid production, whereas pH 7.0, although still effective, resulted in relatively lower lipid content (**Figure 2 B**). The lipid productivities and yields from all four different pH-controlled conditions peaked at around 48 hours though the values at pH 5.0 appeared higher in the middle of the run (**Figure 2 C** and **D**).

**Figure 2.**
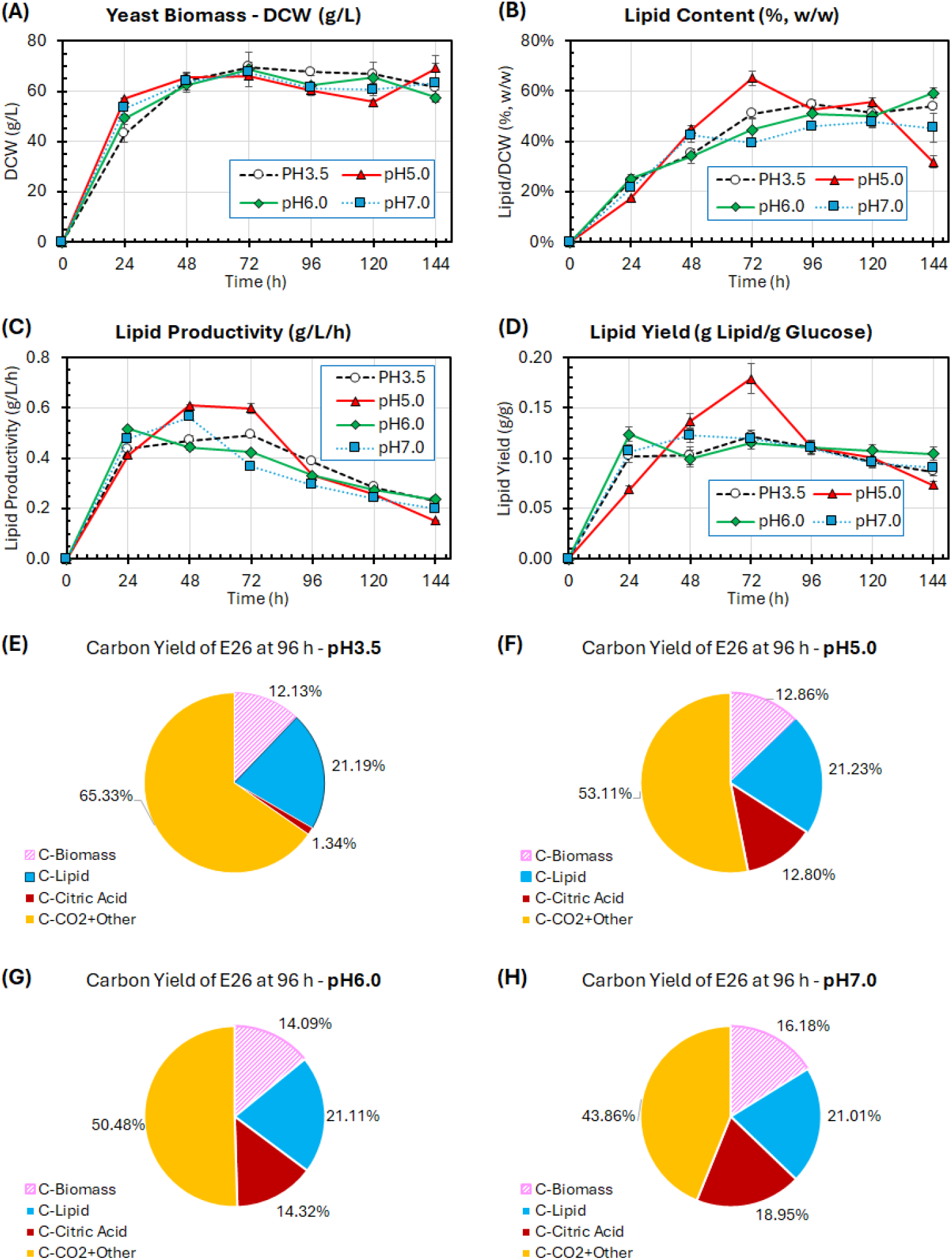
Fed-batch fermentations of *Y. lipolytica* E26 strain with different pH values. (A) Biomass – DCW (g/L), (B) Lipid content - Lipid/DCW (%, w/w), (C) Lipid productivity (g/L/h), and (D) Lipid yield from glucose (g/g). E), (F), (G), and (H) show the molecular carbon conversion yield from glucose into lipid-free biomass, lipid, citric acid, and CO_2_ for pH 3.5, 5.0, 6.0, and 7.0, respectively. The error bars represent the standard deviation of two independent replicates.

It is well-known that the *Y. lipolytica* can tolerate lower pH conditions as compared with most other microorganisms^20^. However, literature underlines that the most suitable pH levels for metabolite production are slightly under neutral pH (5.5-6.0)^21,22^. Our fed-batch fermentation results were consistent with the previous findings on the pH effect, but we also found that a lower pH value such as 3.5 may still be feasible for large scale application, especially an acidic environment may minimize the need of using base for pH control and reduce the risk of bacterial contamination as most bacteria cannot tolerate low pH levels. Interestingly, further analysis on carbon yield of the all fermentation products (lipid-free biomass, lipid, citric acid, and CO_2_ + other byproducts) showed that the carbon yield of the main byproduct citric acid (1.3%, C in citric acid/total C in glucose) was significantly lower than those (13∼19%) from the pH 5.0∼7.0 (**Figure 2 E-H**). However, carbon loss as CO_2_ from pH3.5 (up to 65%) was significantly higher than from the pH 5.0∼7.0 (44∼53%), indicating a complicated mechanism inside the yeast cells to assimilate glucose into more CO_2_ for enhanced energy generation and regulations under an acidic environment^23,24^. The carbon yield of lipid-biomass increased from 12% to 16% as the pH changed from 3.5 to 7.0, indicating a neutral environment is preferred for biomass formation. In addition, the carbon yields of lipid for all four different pH conditions were similar, around 21% (**Figure 2 E-H**). Overall, the pH5∼6 with relatively higher lipid production rate and yield was suggested for further investigation under CF conditions.

### 3.2. Model simulation for continuous fermentation with different scenarios

It is labor-intensive to use experiments to determine the optimal condition for a CF process as each CF experiment takes weeks of time. In this study, a kinetic model for *Y. lipolytica* fermentation was used to simulate a series of feed (glucose, yeast extract, ammonium sulfate) and operating conditions (feeding rate and dilution rate), which helped guide the design of a new CF in the 1-L bioreactor experiment. The model was derived from our previous study of using *Y. lipolytica* for lipid and omega-3 fatty acid production^4^, which was further modified for producing up to 80% lipid in the yeast biomass, as described in the **Supplementary Information**. The model was used to simulate 36 different scenarios to achieve > 100 g/L biomass, > 60% lipid in DCW, > 0.2 g/g lipid yield, and nearly 2 g/L/h lipid productivity within an oxygen uptake rate (OUR) capacity of 120 mmol/L/h, which is achievable in the 1-L bioreactor system with a maximum stirring speed of 1200 rpm (**Table 1**). After careful consideration, Case 21 was selected to design the initial feed and dilution rate for the CF experiment: dilution rate of 0.25 h^-1^ (or residence time of 40 h) with the feed consisting of 360 g/L glucose, 16 g/L yeast extract, and 12 g/L (NH_4_)_2_SO_4_. Other conditions include pH 6 and dissolved oxygen of 20% air saturation^14^. It predicted biomass of 113 g/L, lipid content 65% in biomass, lipid productivity of 1.8 g/L/h, and lipid yield of 0.22 g/g with an ideal strain and under well-controlled, steady-state CF conditions.

**Table 1.**
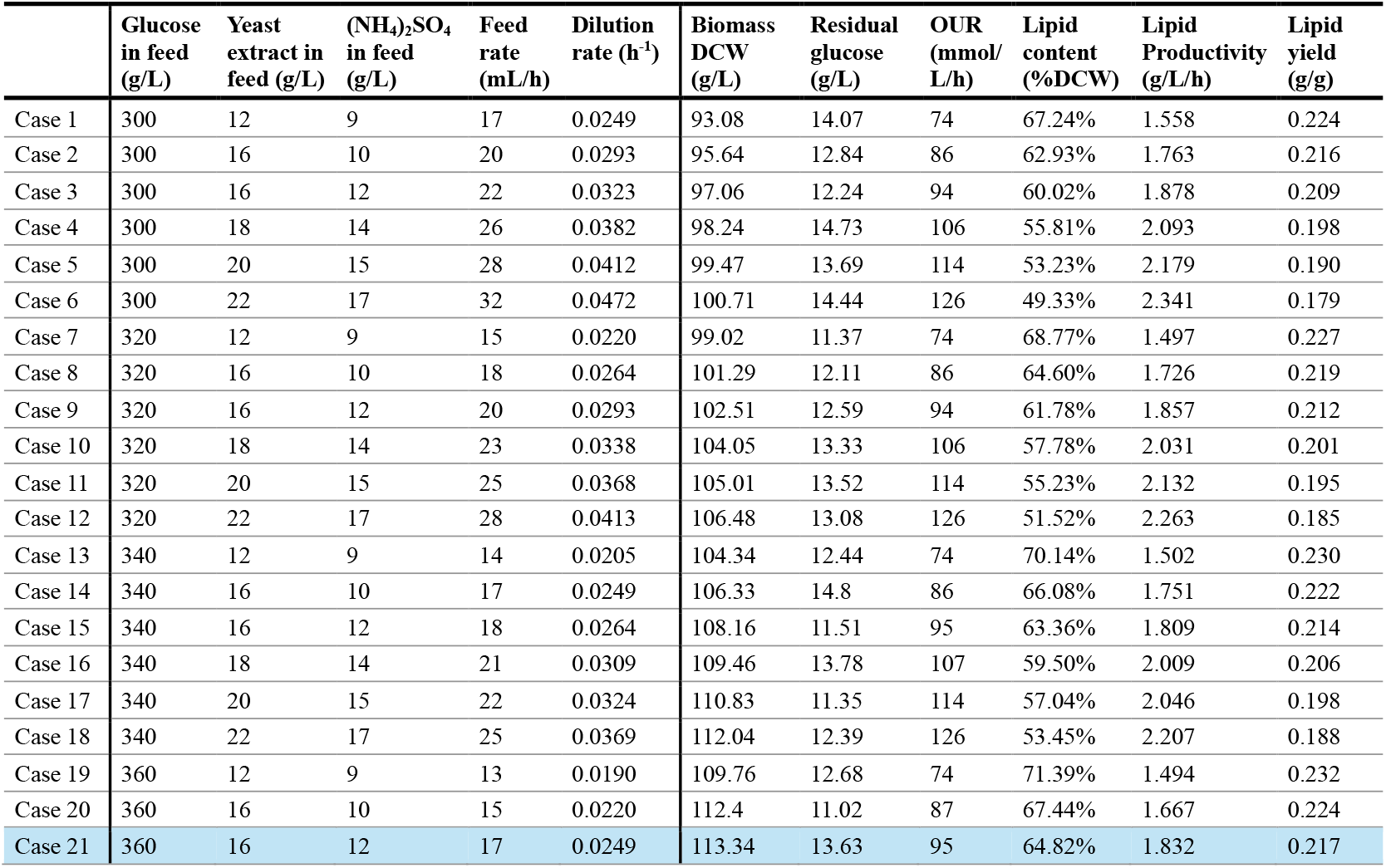
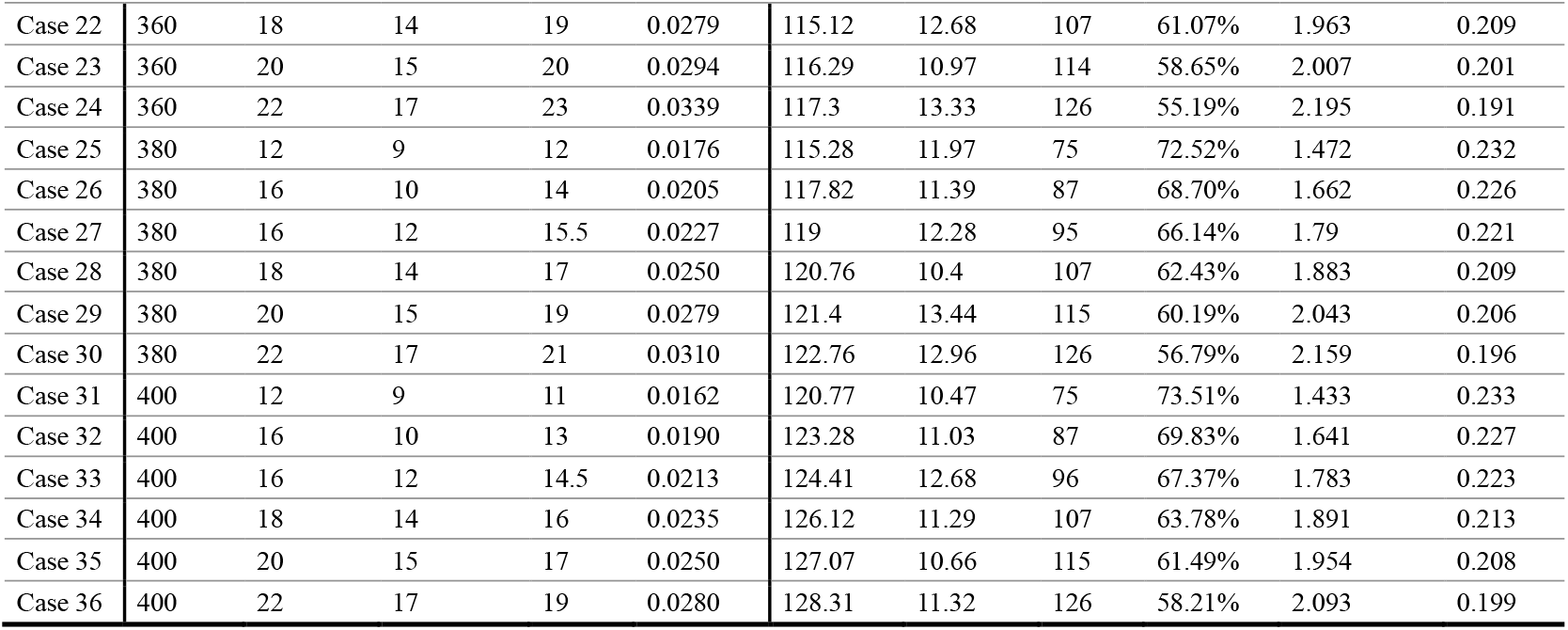
Model simulation results for different scenarios of continuous fermentation in a 1-L bioreactor.

|  | Glucose in feed (g/L) | Yeast extract in feed (g/L) | (NH<sub>4</sub>)<sub>2</sub>SO<sub>4</sub> in feed (g/L) | Feed rate (mL/h) | Dilution rate (h<sup>-1</sup>) | Biomass DCW (g/L) | Residual glucose (g/L) | OUR (mmol/L/h) | Lipid content (%DCW) | Lipid Productivity (g/L/h) | Lipid yield (g/g) |
| --- | --- | --- | --- | --- | --- | --- | --- | --- | --- | --- | --- |
| Case 1 | 300 | 12 | 9 | 17 | 0.0249 | 93.08 | 14.07 | 74 | 67.24% | 1.558 | 0.224 |
| Case 2 | 300 | 16 | 10 | 20 | 0.0293 | 95.64 | 12.84 | 86 | 62.93% | 1.763 | 0.216 |
| Case 3 | 300 | 16 | 12 | 22 | 0.0323 | 97.06 | 12.24 | 94 | 60.02% | 1.878 | 0.209 |
| Case 4 | 300 | 18 | 14 | 26 | 0.0382 | 98.24 | 14.73 | 106 | 55.81% | 2.093 | 0.198 |
| Case 5 | 300 | 20 | 15 | 28 | 0.0412 | 99.47 | 13.69 | 114 | 53.23% | 2.179 | 0.190 |
| Case 6 | 300 | 22 | 17 | 32 | 0.0472 | 100.71 | 14.44 | 126 | 49.33% | 2.341 | 0.179 |
| Case 7 | 320 | 12 | 9 | 15 | 0.0220 | 99.02 | 11.37 | 74 | 68.77% | 1.497 | 0.227 |
| Case 8 | 320 | 16 | 10 | 18 | 0.0264 | 101.29 | 12.11 | 86 | 64.60% | 1.726 | 0.219 |
| Case 9 | 320 | 16 | 12 | 20 | 0.0293 | 102.51 | 12.59 | 94 | 61.78% | 1.857 | 0.212 |
| Case 10 | 320 | 18 | 14 | 23 | 0.0338 | 104.05 | 13.33 | 106 | 57.78% | 2.031 | 0.201 |
| Case 11 | 320 | 20 | 15 | 25 | 0.0368 | 105.01 | 13.52 | 114 | 55.23% | 2.132 | 0.195 |
| Case 12 | 320 | 22 | 17 | 28 | 0.0413 | 106.48 | 13.08 | 126 | 51.52% | 2.263 | 0.185 |
| Case 13 | 340 | 12 | 9 | 14 | 0.0205 | 104.34 | 12.44 | 74 | 70.14% | 1.502 | 0.230 |
| Case 14 | 340 | 16 | 10 | 17 | 0.0249 | 106.33 | 14.8 | 86 | 66.08% | 1.751 | 0.222 |
| Case 15 | 340 | 16 | 12 | 18 | 0.0264 | 108.16 | 11.51 | 95 | 63.36% | 1.809 | 0.214 |
| Case 16 | 340 | 18 | 14 | 21 | 0.0309 | 109.46 | 13.78 | 107 | 59.50% | 2.009 | 0.206 |
| Case 17 | 340 | 20 | 15 | 22 | 0.0324 | 110.83 | 11.35 | 114 | 57.04% | 2.046 | 0.198 |
| Case 18 | 340 | 22 | 17 | 25 | 0.0369 | 112.04 | 12.39 | 126 | 53.45% | 2.207 | 0.188 |
| Case 19 | 360 | 12 | 9 | 13 | 0.0190 | 109.76 | 12.68 | 74 | 71.39% | 1.494 | 0.232 |
| Case 20 | 360 | 16 | 10 | 15 | 0.0220 | 112.4 | 11.02 | 87 | 67.44% | 1.667 | 0.224 |
| Case 21 | 360 | 16 | 12 | 17 | 0.0249 | 113.34 | 13.63 | 95 | 64.82% | 1.832 | 0.217 |
| Case 22 | 360 | 18 | 14 | 19 | 0.0279 | 115.12 | 12.68 | 107 | 61.07% | 1.963 | 0.209 |
| Case 23 | 360 | 20 | 15 | 20 | 0.0294 | 116.29 | 10.97 | 114 | 58.65% | 2.007 | 0.201 |
| Case 24 | 360 | 22 | 17 | 23 | 0.0339 | 117.3 | 13.33 | 126 | 55.19% | 2.195 | 0.191 |
| Case 25 | 380 | 12 | 9 | 12 | 0.0176 | 115.28 | 11.97 | 75 | 72.52% | 1.472 | 0.232 |
| Case 26 | 380 | 16 | 10 | 14 | 0.0205 | 117.82 | 11.39 | 87 | 68.70% | 1.662 | 0.226 |
| Case 27 | 380 | 16 | 12 | 15.5 | 0.0227 | 119 | 12.28 | 95 | 66.14% | 1.79 | 0.221 |
| Case 28 | 380 | 18 | 14 | 17 | 0.0250 | 120.76 | 10.4 | 107 | 62.43% | 1.883 | 0.209 |
| Case 29 | 380 | 20 | 15 | 19 | 0.0279 | 121.4 | 13.44 | 115 | 60.19% | 2.043 | 0.206 |
| Case 30 | 380 | 22 | 17 | 21 | 0.0310 | 122.76 | 12.96 | 126 | 56.79% | 2.159 | 0.196 |
| Case 31 | 400 | 12 | 9 | 11 | 0.0162 | 120.77 | 10.47 | 75 | 73.51% | 1.433 | 0.233 |
| Case 32 | 400 | 16 | 10 | 13 | 0.0190 | 123.28 | 11.03 | 87 | 69.83% | 1.641 | 0.227 |
| Case 33 | 400 | 16 | 12 | 14.5 | 0.0213 | 124.41 | 12.68 | 96 | 67.37% | 1.783 | 0.223 |
| Case 34 | 400 | 18 | 14 | 16 | 0.0235 | 126.12 | 11.29 | 107 | 63.78% | 1.891 | 0.213 |
| Case 35 | 400 | 20 | 15 | 17 | 0.0250 | 127.07 | 10.66 | 115 | 61.49% | 1.954 | 0.208 |
| Case 36 | 400 | 22 | 17 | 19 | 0.0280 | 128.31 | 11.32 | 126 | 58.21% | 2.093 | 0.199 |

### 3.3. Single-stage continuous fermentation

The single-stage CF experiment started with the dilution rate and feed conditions as shown in Case 21 of the simulation results (**Table 1**). To investigate the effects of major process parameters on the fermentation performance, the following process changes were applied during the entire CF period, which include (1) increasing the pH value from 6.0 to 7.0 at 216 h, (2) lowering the pH value from 7.0 to 5.0 at 360 h, (3) increasing the feeding glucose concentration from 360 g/L to 460 g/L at 456 h, (4) lowering the DO change from 20% to 10% air saturation at 744 h, (5) changing all parameters back to their original settings (glucose in feed 360 g/L, pH 6.0, DO 20% air saturation) at 960 h, and (6) lowering the PH value from 6.0 to 3.5 at 1080 h. The dilution rate was maintained at 0.025∼0.03 h^-1^ to maintain the residual glucose within 10∼20 g/L.

As shown in **Figure 3A**, the yeast biomass reached around 100 g/L after 168 h and was maintained at the density until 500 h. After that, it started to gradually increase to > 130 g/L, much higher than what was predicted by model (113 g/L). This may be partially caused by increasing the glucose feed concentration from 360 g/L to 460 g/L at 456 h, which significantly reduced the dilution rate from 0.025∼0.3 h^-1^ to 0.21 h^-1^ and increased the residence time for biomass buildup. Interestingly, changing the glucose feed concentration back to 360 g/L at 960 h did not lead to the decrease of biomass back to 100 g/L. Instead, about 120 g/L DCW was still observed at ∼ 1080 h, which suggests certain mutations were induced and certain cells already evolved to growth faster and generate more biomass. This can also be seen from the increase in the carbon yield of lipid-free biomass from 22% at 96 h to 37% at 1008 h (**Figure 3 E** and **F**).

**Figure 3.**
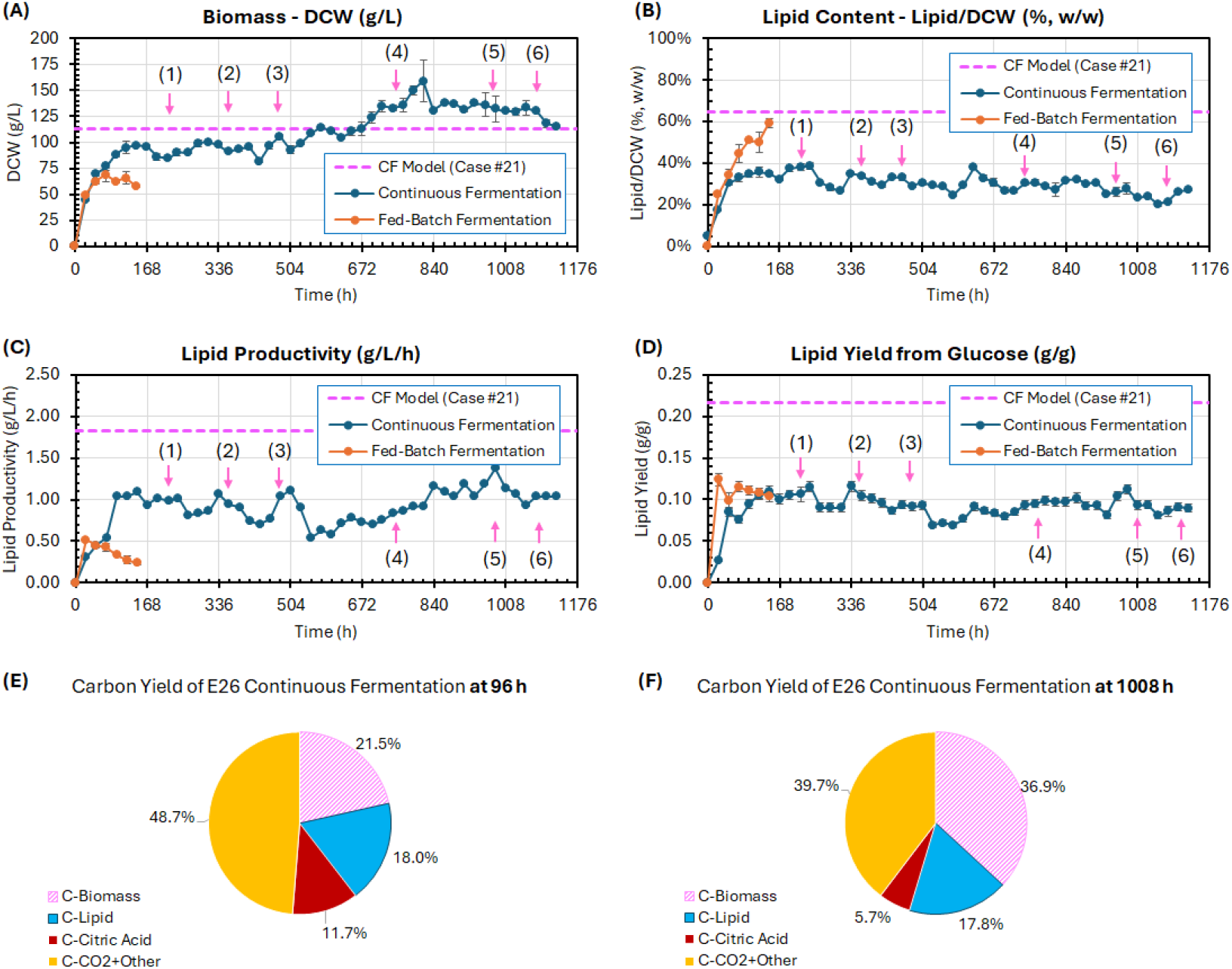
Comparison between the single-stage CF and fed-batch fermentation with *Y. lipolytica* E26 strain. (A) Biomass (g DCW/L), (B) Lipid content in biomass - Lipid/CDW (%, w/w), (C) Lipid productivity (g/L/h), and (D) Lipid yield from glucose (g/g). Numbers with arrows indicate the process changes made to the CF: (1) increasing the pH value from 6.0 to 7.0 at 216 h, (2) lowering the pH value from 7.0 to 5.0 at 360 h, (3) increasing the feeding glucose concentration from 360 g/L to 460 g/L at 456 h, (4) lowering the DO change from 20% to 10% air saturation at 744 h, (5) changing all parameters back to their original settings (glucose in feed 360 g/L, pH 6.0, DO 20% air saturation) at 960 h, and (6) lowering the PH value from 6.0 to 3.5 at 1080 h. Red dashed line shows the CF model predicted level (Case #21, Table 1). (E) and (F) show the carbon yield from glucose into lipid-free biomass, lipid, citric acid, and CO_2_ for the CF at 96 h and 1008 h, respectively. The error bars represent the standard deviation of duplicate measurements.

Lipid is the main product of the *Y. lipolytica* yeast in this study. Lipid content as a percentage of biomass (%, lipid/DCW) slowly dropped from the initially 40% to 30 % during the entire CF process (**Figure 3B**), partially caused by the slow increase in DCW from ∼ 100 g/L to ∼130 g/L (**Figure 3B**). The lipid productivity and yield remained similar at ∼ 1 g/L/h and ∼0.1 g/g, respectively (**Figure 3C** and **3D**). Overall, all actual lipid content, productivity, and yield from the CF experiment were lower than what the model predicted for the initial CF settings. The main reason is that the model parameters in this study were determined by the literature reported lipid production capability of up to 80% lipid content in DCW for fed-batch fermentation for E26^15^, which was significantly higher than what we observed in 1-L fed-batch experiment in this study (40∼60%, as seen in **Figure 2**). Though it is expected that further calibration of the model parameters with additional fed-batch and CF experiments should be able to improve the accuracy of model prediction, the current model still provided good guidance for design of the initial setting of the feed and dilution rate of the CF experiment in a 1-L bioreactor within its achievable OUR (120 mmol/L/h), which successfully helped avoid the need of a number of CF experiments to predetermine these conditions.

In terms of the effects of major process parameters in CF for lipid production, it seems the glucose concentration in feed impacted the fermentation the most, followed by DO and pH value. Higher glucose concentration in feed with proportional increase in nitrogen source (yeast extract and/or (NH_4_)_2_SO_4_) is beneficial to higher cell density and overall productivity, but it may require higher oxygen transfer rate (OTR) capacity of the bioreactor, which is typically subject to a limit of 100∼150 mmol/L/h for most industrial scale bioreactors. Oxygen is important for *Y. lipolytica* cell growth and lipid production. A decrease in DO from 20% to 10% air saturation at 744 h slightly reduced biomass titer. Though a higher DO is preferred for enhanced biomass and lipid production, increased cell density may increase the broth viscosity and reduce the oxygen permeability to the cells^25^. On the other hand, a lower DO may help activate an alternative mitochondrial electron transport chain for energy supply to promote biomass accumulation^26^. Overall, a DO at ∼20% air saturation is appropriate for cell growth and lipid production in CF. No significant effect was observed when the pH changed within 5.0∼7.0, but a significantly loss of biomass titer was seen when the pH value was lowered from 5.0 to 3.5 at 1008 h. Since a relatively lower pH value may reduce the risk of bacterial contamination in a long-term CF period, it is suggested that a pH value near 5.0 should be considered for CF at larger scale.

Further analysis of the carbon yield of major fermentation products, including lipid-free biomass, lipid, citric acid, and others (mainly CO_2_) showed that cells evolved significantly after 1008 h (six weeks) with biomass carbon yield increased from 22% to 36% and citric acid carbon yield decreased from 12% to 6% (**Figure 3E** and **F**). Interestingly, the lipid carbon yield remained at ∼18% throughout the entire CF period. The results suggests that certain mutants were generated with higher lipid-production and lower citric acid production after ALE in the high-cell-density continuous bioreactor, which provides us an opportunity to isolate better lipid producers from the end of the continuous bioreactor. ALE has been widely used to isolate mutants with enhanced cell growth and/or production by achieving increased tolerance of stressful conditions ^27,28^. Typically, it is performed through a series of passages of flask cultures for up to months with cell growth for hundreds of generations. Our study is among the very few using ALE in continuous bioreactor to isolate improved mutants.

Compared with the fed-batch fermentation, the single-stage CF in study improved the overall lipid productivity from 0.3∼0.5 g/L/h to ∼1 g/L/h and maintained the lipid yield at ∼ 0.1 g/g. While cell growth and lipid production prefer different optimal condition, the single-sage CF in this study integrates both cell growth and lipid production in the same bioreactor, which means only sub-optimal conditions can be achieved. Therefore, a two-stage^4^ or even muti-stage CF^29,30^ should be considered in future to further improve the overall lipid titer, rate, and yield, where cell growth and lipid production can be decoupled and carried out in separate bioreactors with each independently controlled at its own optimal conditions to achieve the overall best fermentation performance.

### 3.4. Screening high-lipid producing mutants from ALE in continuous bioreactor

As the ALE in continuous bioreactor generated mutants with potentially higher lipid-production capabilities, the sample collected at 1008 h from the CF experiment was diluted by 10^7^ fold, plated on YNB plate to generate single colonies for screening. Fifteen colonies were randomly picked and tested for further culture in shake flasks (**Figure 1**). The end-of-run samples from the flask experiments were used for biomass and lipid content analysis, as shown in **Figure 4**. Colony #3, #6, and # 12 exhibited higher biomass production than the original E26 (12 g/L), with #6 producing the highest, surpassing 14 g/L, demonstrating a clear adaptive advantage in terms of growth under the tested conditions. For lipid production, both Colony #3 and #12 achieved > 60% lipid/DCW while most other colonies had lipid contents within 20∼50%. It seemed the variation in lipid content between different colonies were significantly larger than that in biomass. The two mutants from colony #3 and #12 with the highest lipid production were renamed as strain E26E03 and E26E12, respectively, which were further tested in 1-L fed-batch fermentation to verify their improved lipid production capability over the original strain E26.

**Figure 4.**
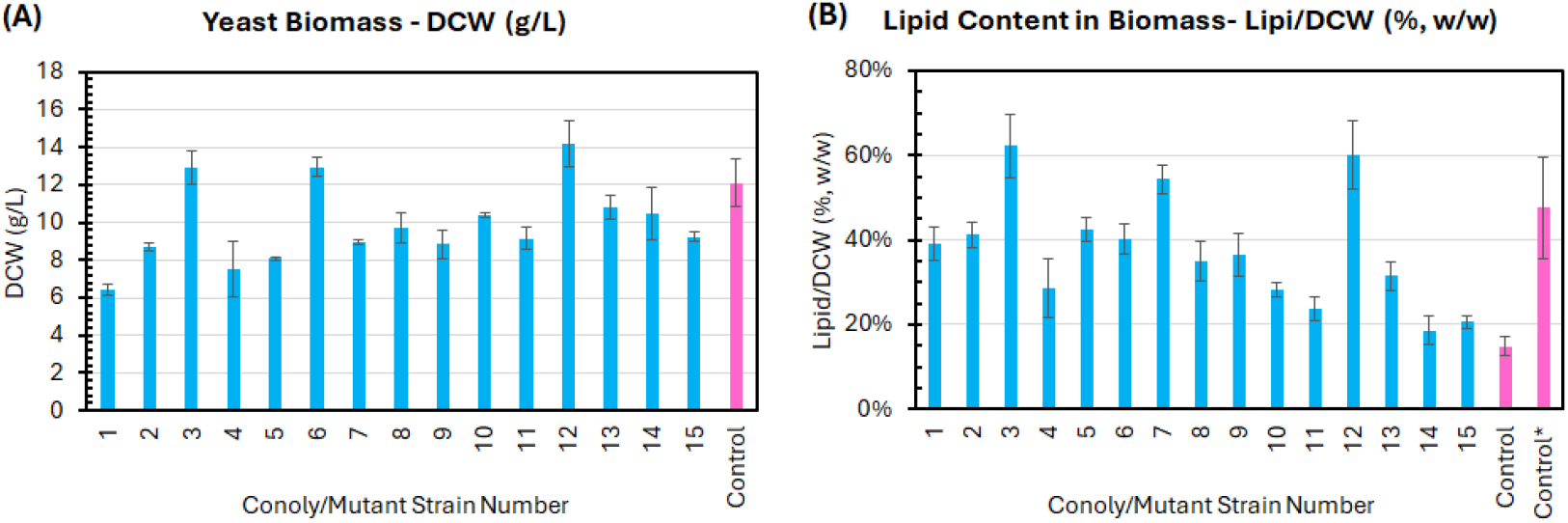
Flask culture screening of the CF mutants. (A) Biomass (g/L) and (B) Lipid/CDW (%). The error bars represent the standard deviation of two independent replicates. The original strain E26 was used as the control to compare with all mutants from the CF experiment. The Control* for lipid content refers to the average lipid data from the end of the run of four fed-batch fermentation experiments.

As seen in **Figure 5**, both mutant strains E26E03 and E26E12 accumulated higher biomass that the original strain E26. The lipid content, productivity, and yield from the two mutants were also significantly higher than from E26, which verified the results from the flask experiments. The best mutant strain E26E03 improved the biomass, lipid content, and yield by 43%, 30%, and 51%, respectively over the original strain E26 at 144 h. This indicates the large population of the CF preserved different mutants with different characteristics, without harming to each other^31^. Further analysis of the carbon yield of all fermentation products for E26E03 showed that very minimal carbon flow went to the byproduct citric acid (2.5%) as compared to E26 (14.3%). This also led to a significant increase in lipid carbon yield from 21% to 28% (**Figure 5E, F**, and **G**). In addition, we noticed that the mutant strain E26E03 grew faster and consumed nearly 60% more glucose than E26 during the fed-batch fermentation, probably caused by a much lower stressful environment due to the much less accumulation of the byproduct citric acid (12.5 g/L vs. 58 g/L).

**Figure 5.**
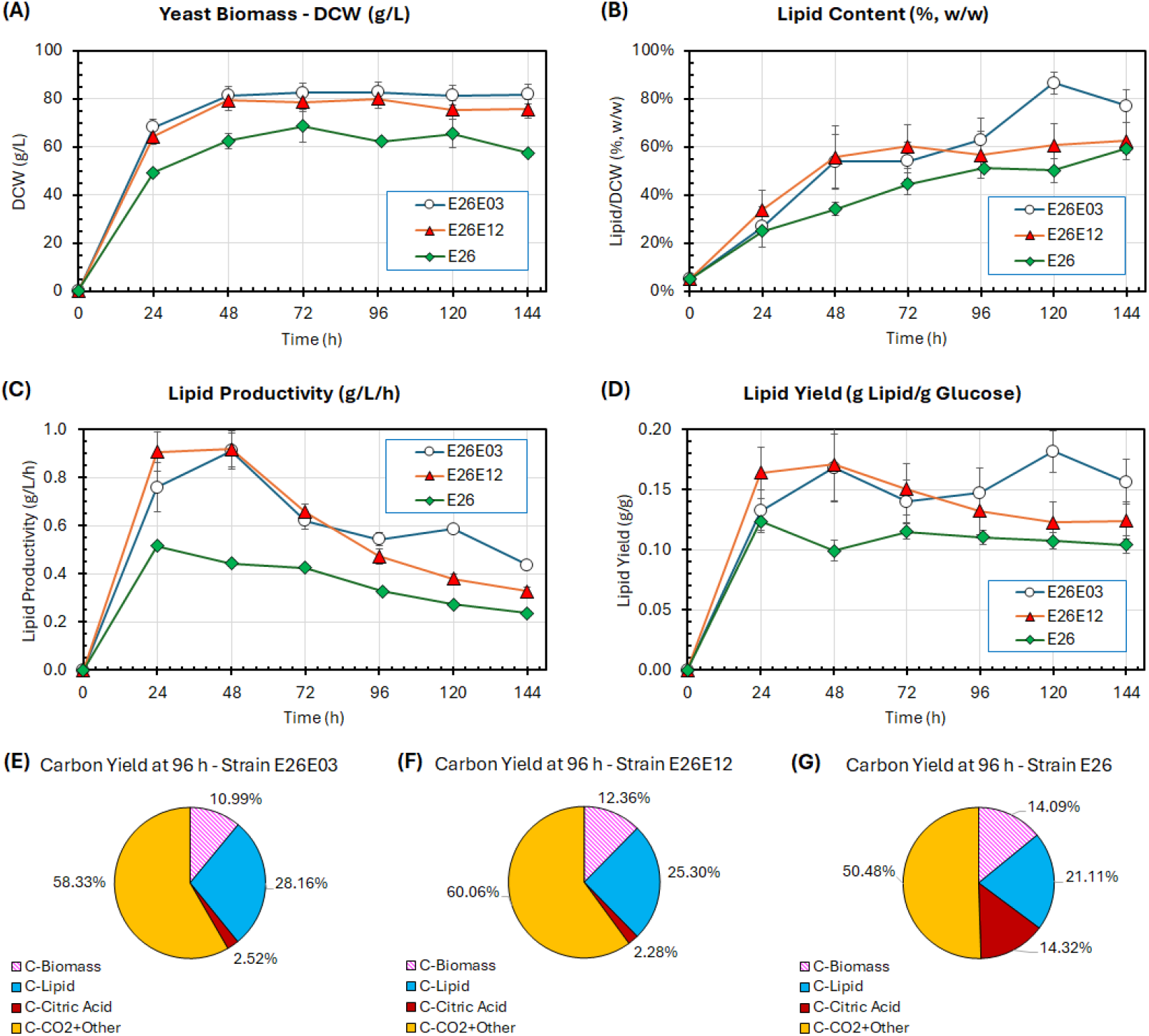
Comparison between the fed-batch fermentation of two selected mutants (E26E03 and E26E12) with the original strain E26. (A) Yeast biomass, (B) Lipid content in biomass – Lipid/CDW (%, w/w), (C) Lipid productivity (g/L/h), (D) Lipid yield from glucose (g/g). (E), (F), and (G) show the molecular carbon conversion yield from glucose into lipid-free biomass, lipid, citric acid, and CO_2_ for strain E26E03, E26E12, and the original strain E26, respectively. The error bars represent the standard deviation of two independent replicates.

## 4. Conclusion

This study demonstrates the potential of integrating model-guided CF with ALE in continuous bioreactor to enhance lipid production by the engineered yeast *Y. lipolytica* E26. Fed-batch fermentation determined the appropriate ranges of pH value for enhanced lipid production and provided basic understandings of the strain and fermentation process, which helped design and optimize CF for further improved lipid productivity. Kinetic model simulation for a series of feed and operating conditions was used to determine the initial process settings for the CF in the 1-L bioreactor experiment. Compared with the fed-batch fermentation process, the single-stage CF successfully improved lipid productivity from 0.3∼0.5 g/L/h to about 1 g/L/h while maintained the lipid yield at around 0.1 g/g. In addition, the long-term CF also generated mutants with significantly faster cell growth, higher lipid production, and less byproduct formation, indicating the feasibility of using ALE in continuous bioreactor to isolate mutant strain with significantly higher lipid production. Our findings highlight the major research progress in integrating predictive model, continuous fermentation, and evolutionary strategy for more sustainable and profitable biomanufacturing. Due to the limitation of single-stage CF that fails to decouple cell growth and production and simultaneously optimize them, two-stage or multi-stage CF should be explored in future to further improve production titer, rate, and yield.

## Author Contributions

EK: Conceptualization, method development, experiments, data analysis, writing – original, writing – review & editing; LS: experiments; EA: experiments; DX: Conceptualization, supervision, funding acquisition, writing – original, writing – review & editing. All authors agree to the final manuscript.

## Acknowledgement

This work was funded by the NSF/CBET (Award Number 2133660). The authors would also like to thank Dr. Hal Alper for providing the *Y. lipolytica* strain E26 for the research. We also like to thank Drs. Carl Lawton and Jin Xu for their support for the use of equipment and facilities in Massachusetts Biomanufacturing Center.

## Supplementary Information

### Kinetic model development for fed-batch and continuous fermentation: Cell growth

The biomass [X] consists of lipid-free cell mass [X_f_] and lipid [L]:

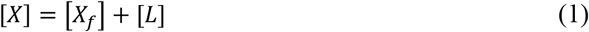

where the unit for [X], [X_f_], and [L] is [g/L]. The cell growth equation can be described with the specific growth rate (μ) (Eq. 2). All concentrations, including [X], [X_f_], and [L], in the culture medium were calibrated by considering the volume change caused by the dilution rate D and evaporation rate *V*_*evap*_(see Eq. 19∼21) during the fed-batch or continuous fermentation process:

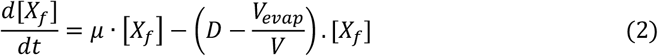

The modified Monod-type equations (Eq. 3 and 5) were used to describe the yeast’s specific growth rate (µ) (1/h) at different glucose (S, g/L), dissolved oxygen (O, air saturation%), and nitrogen (N, g/L) concentrations. When nitrogen is not limited, cells grow through using the extracellular nitrogen (provided by ammonium sulfate and yeast extract), which is described by Eq. 4. The Y_X/N_/10 is the saturation constant for cell growth on intracellular nitrogen (*N*_*cell*_) and was determined by empirical data. When all available extracellular nitrogen is consumed, cells continue to grow at a certain rate, expressed by Eq. 5.

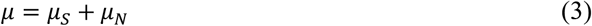

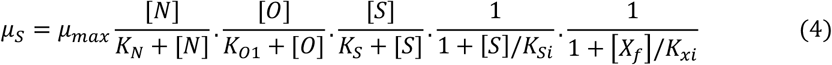

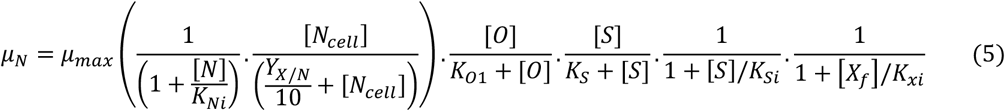

where all the symbols are described below:

D dilution rate (1/h)

N nitrogen concentration (mM) S glucose concentration (g/L)

*K*_*N*_ saturation constant for intracellular nitrogen in growth kinetics (g/L)

*K*_*O1*_ saturation constant for dissolved oxygen in growth & glucose uptake kinetics (% Air)

*K*_*S*_ saturation constant for glucose utilization (g/L)

*K*_*Si*_ inhibition constant for glucose in lipid-based growth kinetics (g/L)

*K*_*Xi*_ constant for cell density effect on cell growth and lipid/byproduct formation (unit/L)

*N*_*cell*_ intracellular stored nitrogen available for cell growth (g/g)

*V* working volume of the bioreactor vessel (L)

*V*_*evap*_ Evaporation rate (L/h)

*K*_*Ni*_ nitrogen limitation constant to trigger on lipid and byproduct production (g/L)

*μ*_*S*_ Specific growth rate when extracellular nitrogen is not limited (1/h)

*μ*_*N*_ Specific growth rate when extracellular nitrogen is limited (1/h)

### Extracellular nitrogen consumption

The total nitrogen consumption rate is described by Eq. (6).

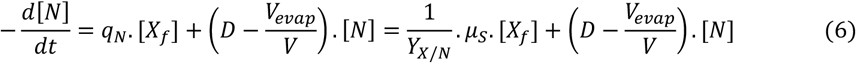

where all new symbols are described below:

N nitrogen concentration (mM)

q_*N*_ Specific nitrogen consumption rate (1/h)

*N*_*cell*_ intracellular stored nitrogen available for cell growth (g/g)

*K*_*Ni*_ nitrogen limitation constant to trigger on lipid and byproduct production (g/L)

*Y*_*X*/*N*_ yield coefficient of cell mass based on nitrogen consumed (g/g)

### Intracellular nitrogen consumption

The intracellular nitrogen content accumulates from 0 to (1/3*YY*_*XX*/*NN*_) when nitrogen concentration [N] in the medium is non-limited and is subsequently utilized for further growth once nitrogen becomes limited. The model assumes that during the growth phase, *N*_*cell*_increases from 0 to 1/3 (1/Y_X/N_) under non-limited nitrogen conditions and gradually depletes to 0 under nitrogen-limited conditions in the oleaginous phase.

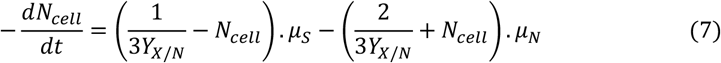

### Byproduct formation

By-product [C] (mainly citric acid) accumulation is also considered and assumed as more agile under nitrogen-limited and glucose-rich conditions. The model posits that only the *r*_*L*_portion of the total carbon flux (byproduct and lipid) in the lipid synthesis pathway contributes to lipid formation due to overflow loss in the byproduct. Additionally, the model incorporates a tolerance limit (K_ci_) for the byproduct-to-cell mass ratio (*C*/*X*_f_) and accounts for the effect of cell density on product formation through a term defined as 1/(1+ *X*_f_/*K*_xi_), as shown in Eq. 8-10.

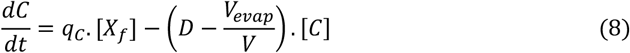

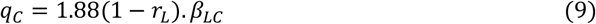

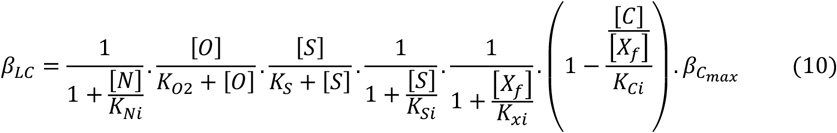

where all new symbols are described below:

q_*C*_ specific byproduct production rate (1/h)

*r*_*L*_ constant ratio of lipid carbon flow to total carbon flow toward citrate and lipid (unitless)

*β*_*LC*_ specific rate of total carbon introduced into lipid and byproduct synthesis (1/h)

*K*_*O2*_ saturation constant for dissolved oxygen in lipid, byproduct, and lipid kinetics (% Air)

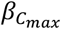 coefficient of maximum byproduct production for cell mass (1/h)

### Lipid accumulation

The lipid production model is formulated using partial growth-associated kinetics. Lipid accumulation in the oleaginous phase occurs under nitrogen-limited, glucose-rich, and aerobic conditions (Eq. 11-12). A fraction of the accumulated lipids may degrade when their concentration reaches high levels in the presence of oxygen (Eq. 13 and 14).

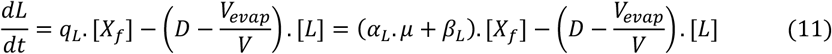

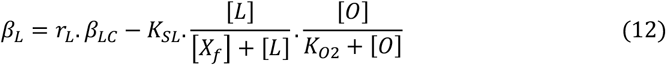

where all new symbols are described below:

q_*L*_specific lipid production rate (1/h)

*α*_*L*_ coefficient of lipid production for cell growth

*β*_*L*_ coefficient of lipid production for cell mass (1/h)

*K*_*SL*_ coefficient for lipid consumption/decomposition (1/h)

### Glucose consumption

The glucose consumption rate is -r_s_, where r_s_ = d[S]/dt and [S] is the glucose concentration. Like specific growth rate, specific substrate (glucose) consumption rate can be calculated as Eq. 13 and Eq. 14, respectively.

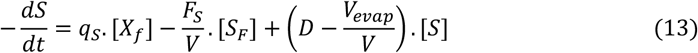

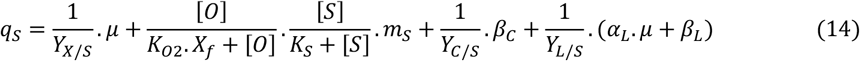

where all new symbols are described below:

*S*_*F*_ glucose concentration in glucose feed (g/L)

*m*_*S*_ maintenance coefficient for glucose (g/g/h)

*Y*_*C*/*S*_ yield coefficient of byproduct based on glucose consumed (g/g)

*Y*_*L*/*S*_ yield coefficient of lipid based on glucose consumed (g/g)

### Dissolved oxygen, oxygen transfer rate (OTR), and oxygen uptake rate (OUR)

The dissolved oxygen (DO, represented by 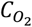 or [O]) in the bioreactor can be described by Eq. 15:

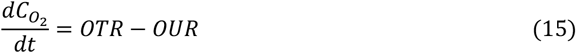

where OTR corresponds to the oxygen transfer rate (unit: mmol/L/h) and OUR (unit: mmol/L/h) is the oxygen uptake rate. Under steady-state control,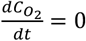. OTR is represented by Eq. 16:

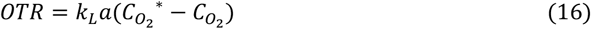

where *k*_*L*_*a* is the volumetric oxygen transfer coefficient (h^-1^), 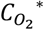 is the saturated DO concentration (mmol/L), and 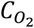 is the actual DO concentration in the broth (mmol/L). The OUR can be calculated for a specific strain by using following empirical equations.

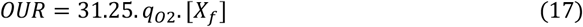

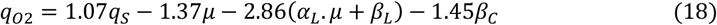

### Reaction volume, dilution, and base feeding

The dilution rate D, the working volume V, and base feed rate can be described by the following equations:

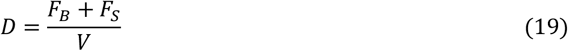

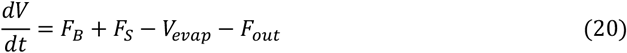

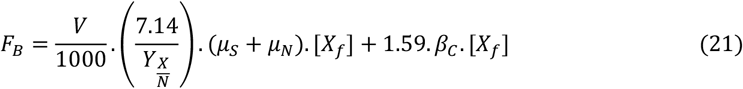

where *F*_B_ is the feed rate of KOH (10 N) for pH control, *F*_S_ is the glucose feed (L/h), *F*_out_ is the rate of fermentation broth withdrawn from the bioreactor (L/h), and *V* is the working volume (L). For fed-batch fermentation, *F*_*out*_=0. For continuous fermentation, *F*_*out*_=*F*_*B*_+*F*_*S*_–*V*_*evap*_.

### Model parameters used in simulation of continuous fermentation

**Table S1.**
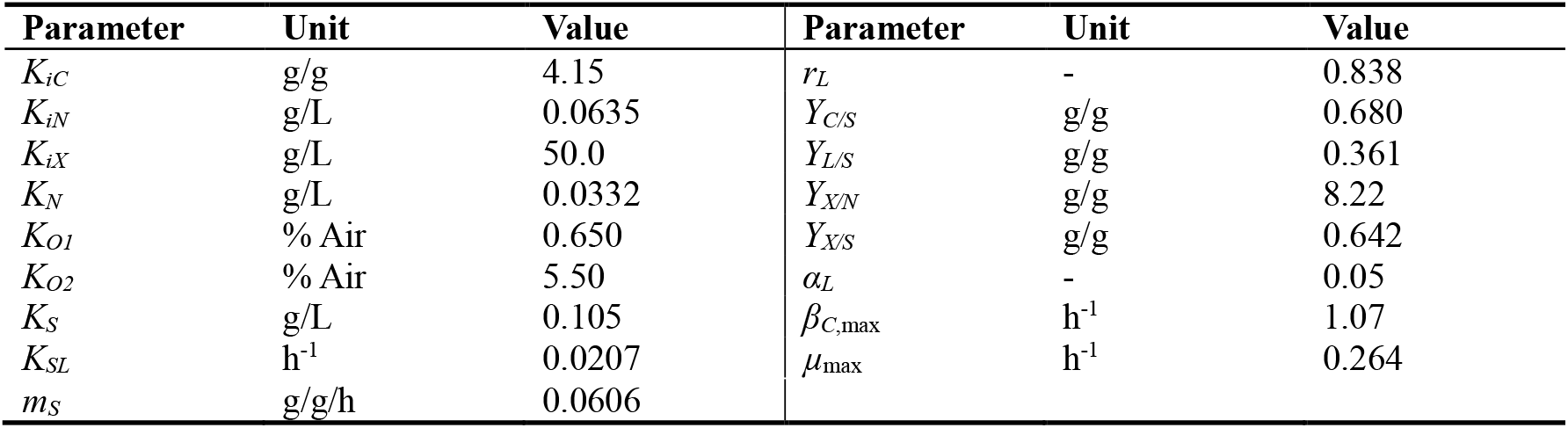
Model parameters used in this study.

| Parameter | Unit | Value | Parameter | Unit | Value |
| --- | --- | --- | --- | --- | --- |
| $K_{iC}$ | g/g | 4.15 | $r_L$ | - | 0.838 |
| $K_{iN}$ | g/L | 0.0635 | $Y_{C/S}$ | g/g | 0.680 |
| $K_{iX}$ | g/L | 50.0 | $Y_{L/S}$ | g/g | 0.361 |
| $K_N$ | g/L | 0.0332 | $Y_{X/N}$ | g/g | 8.22 |
| $K_{O1}$ | % Air | 0.650 | $Y_{X/S}$ | g/g | 0.642 |
| $K_{O2}$ | % Air | 5.50 | $\alpha_L$ | - | 0.05 |
| $K_S$ | g/L | 0.105 | $\beta_{C,\max}$ | h<sup>-1</sup> | 1.07 |
| $K_{SL}$ | h<sup>-1</sup> | 0.0207 | $\mu_{\max}$ | h<sup>-1</sup> | 0.264 |
| $m_S$ | g/g/h | 0.0606 | | | |

